# Dietary methionine/choline deficiency affects behavioural and molecular circadian rhythms

**DOI:** 10.1101/2024.01.04.574014

**Authors:** Benjamin Saer, George Taylor, Andrew Hayes, Bharath Ananthasubramaniam, Jean-Michel Fustin

**Affiliations:** The University of Manchester; Faculty of Biology, Medicine and Health; Centre for Biological Timing, The Meth Lab; The University of Manchester; Faculty of Biology, Medicine and Health; Genomic Technologies Core facility; The University of Manchester; Faculty of Biology, Medicine and Health; Biological Mass Spectrometry Facility; The Humboldt University of Berlin; Institute for Theoretical Biology

## Abstract

The feeding/fasting cycles controlled by our circadian clock impose great daily metabolic and physiological changes, and yet investigations into the consequences of metabolic deficiencies, either dietary or genetic, have often ignored the time of day or the circadian time of the animals or subjects. In addition, these deficiencies may themselves disrupt our circadian clock, causing secondary metabolic, physiological and behavioural disorders.

Dietary methionine/choline deficiency in rodents is a common model for human non-alcoholic steatohepatitis, but methionine and choline are nutrients essential for many other processes beyond fatty acid synthesis in the liver, including biological methylations and 1-carbon metabolism, regulation of translation notably via the mTOR pathway, phospholipid synthesis, polyamine pathway and glutathione synthesis.

We have previously shown that circadian rhythms in many organisms are highly sensitive to deficiency or excesses of 1-carbon metabolites. Using a methionine/choline deficient diet in mice, we illustrate the nutrigenomic crosstalk between circadian rhythms and 1-carbon metabolism. We show not only that circadian locomotor activity behaviour is profoundly, rapidly and reversibly affected by methionine/choline deficiency, but also that the effects of methionine/choline deficiency on gene expression and 1-carbon metabolites are dependent on circadian time, illustrating the importance of considering circadian rhythms in metabolic studies. This study also highlights the impact of what we eat, or don’t, on our behaviour and biological rhythms.

## Introduction

The methionine/choline deficient (MCD) diet is a widespread method to induce hepatic steatosis in rodent as a model for human non-alcoholic steatohepatitis (NASH)^1^. The mechanisms leading to steatosis were first attributed to diminished very-low-density lipoprotein (VLDL) assembly and secretion, leading to reduced triglyceride clearance^2^. It was later observed that MCD also caused a reduction in mitochondrial β-oxidation, increased oxidative stress and changes in cytokines and adipokines production leading to steatohepatitis, inflammation and fibrosis^3–7^. The effects of the MCD diet had been shown to be dependent and the age, sex, species and strain of the animal^8^.

While NASH in human is usually associated with obesity and insulin resistance, rodents provided with an MCD diet display severe weight loss and improved insulin sensitivity, questioning the validity of the MCD model^9,10^.

Choline deficiency alone causes steatohepatitis, triglycerides and free fatty acids accumulation in the liver but no weight loss^3^. In contrast, methionine deficiency alone causes weight loss, injury, oxidative stress, fibrosis and inflammation, with somewhat milder accumulation of hepatic triglyceride and free fatty acids compared to CD and MCD diets^3^. These effects of MD and MCD on steatosis and inflammation are likely due to the disruption of methylation-dependant processes, notably phospholipid metabolism critical for hepatic lipid synthesis, such as the trimethylation of phosphatidylethanolamine to phosphatidylcholine^11^. This is largely supported by the hepatic phenotype of mice deficient in enzymes involved in 1-carbon metabolism and in human patients with deficiencies in these enzymes^12^.

The synthesis of the universal methyl donor S-adenosyl-L-methionine (SAM) requires methionine, and MD or MCD diets leads to a rapid decrease in SAM in the liver^3^, as observed *in vitro* with culture medium without methionine^13^. Donating its methyl, SAM becomes S-adenosyl-L-homocysteine (SAH) then is hydrolysed to homocysteine and adenosine^14^. Homocysteine can be remethylated back to methionine using either methyltetrahydrofolate in many tissues or betaine specifically in the liver^15^. In the liver, choline is the precursor for betaine. Further supporting the hypothesis that the MCD-induced steatohepatitis is dependent on methyl metabolism is the demonstration that betaine supplementation in rats fed MCD for 8 weeks could prevent steatohepatitis, although quantification of SAM and SAH in the liver of these animal after an overnight 8h fasting showed betaine supplementation actually decreased SAM and increased SAH compared to the MCD diet alone^16^, preventing firm conclusions to be drawn.

We have previously shown that circadian rhythms that rely on transcription-translation feedback loops are particularly sensitive to pharmacological or dietary interventions targeting the methyl cycle, and that many mRNAs coding for enzymes involved in 1-carbon metabolism have circadian rhythms^13,17–19^. Together with the fact that NASH and liver fibrosis, both conditions driven by the MCD diet, are exacerbated by circadian misalignments^20^, this intimate link between methyl metabolism and circadian rhythms indicate that the MCD diet may also affect the circadian rhythms of the animal, which perhaps contribute to the pathologies observed in this model. Moreover, very few studies have investigated the systemic effects of the MCD diet but, considering the role of the liver as a metabolic hub, a hepatic deficiency in labile methyl groups is bound to affect other tissues as well. In addition, given the widespread rhythms in the expression of enzymes involved in 1-carbon metabolism, the metabolic consequences of the MCD diet should be assessed taking into account the time of day or the endogenous circadian time of the animal, rather than typically, and for convenience for the researcher, during the day when nocturnal rodents are fasting and inactive^3^.

Here we investigated the effects of methionine/choline deficiency on circadian locomotor activity rhythms and underlying molecular oscillations in gene expression and 1-carbon metabolome of the liver and the suprachiasmatic nucleus of the hypothalamus (SCN, where the mammalian master circadian clock is located). Our results show that the MCD diet has a profound effect on circadian behaviour, demonstrating that methionine and choline deficiency disrupts far more than fatty acid metabolism in the liver. At the transcriptional level, MCD causes a widespread reprogramming of the circadian transcriptome in the liver and the SCN. In the SCN, overall changes in the circadian transcriptome are milder than in the liver but a small subset of genes involved in methyl metabolism and circadian behaviour are strongly regulated. At the 1-carbon metabolome level, MCD decreases most hepatic 1-carbon metabolites compared to control fed animals but in a time-dependent manner. In the SCN, pronounced time-dependent differences in the effects of the MCD diet were seen for most 1-carbon metabolites.

Together our results demonstrate that the MCD diet does affect systemic methyl metabolism, associated with circadian rhythms disruption, and illustrate the need to consider circadian time when the metabolic effects of dietary deficiencies are being investigated.

## Results

### Methionine/Choline deficiency disrupts circadian locomotor activity rhythms

We first sought to define the impact of MCD on circadian locomotor activity behaviour of mice. Mice were provided with control or MCD diet and kept for 10 days in standard light/dark cycles for acclimation, after which mice were kept under constant darkness for several weeks to allow the expression of endogenous rhythms. To minimise the impact of the MCD diet on steatosis, our MCD or control diets contained 4% fat (9.1 % kcal from fat) instead of the high 8-10% (18.3-22.1 % kcal from fat) found in standard MCD diets such as Envigo’s TD.90262 or TD.01037.

Surprisingly, the MCD diet had pronounced effects on circadian behaviour in both male and female mice: it shortened the period of locomotor activity rhythms (Fig. 1A, B), caused irregular activity onsets and compressed the active phase to a single intense bout of wheel-running activity at the beginning of the subjective night, a pattern that worsened with time (Fig. 1B, C). Virtually no activity was detected in the end of the subjective night/early morning, when a bout of activity is expected in control mice (Fig. 1B, C). As expected, the MCD diet caused progressive weight loss (Fig. 1D), but there was no significant differences in food intake between the two groups (Fig. 1E). The effects of MCD on circadian rhythms were however nearly instantaneous, already obvious on the second day under the MCD diet (Fig. 1F), indicating the effects of circadian rhythms are not dependent on chronic starvation or weight loss and are not a secondary consequences of chronic steatosis. Just as quickly, mice immediately recovered from the effects of the MCD diet on circadian rhythms one day after being presented with the control diet (Fig. 1G).

**Figure 1:**
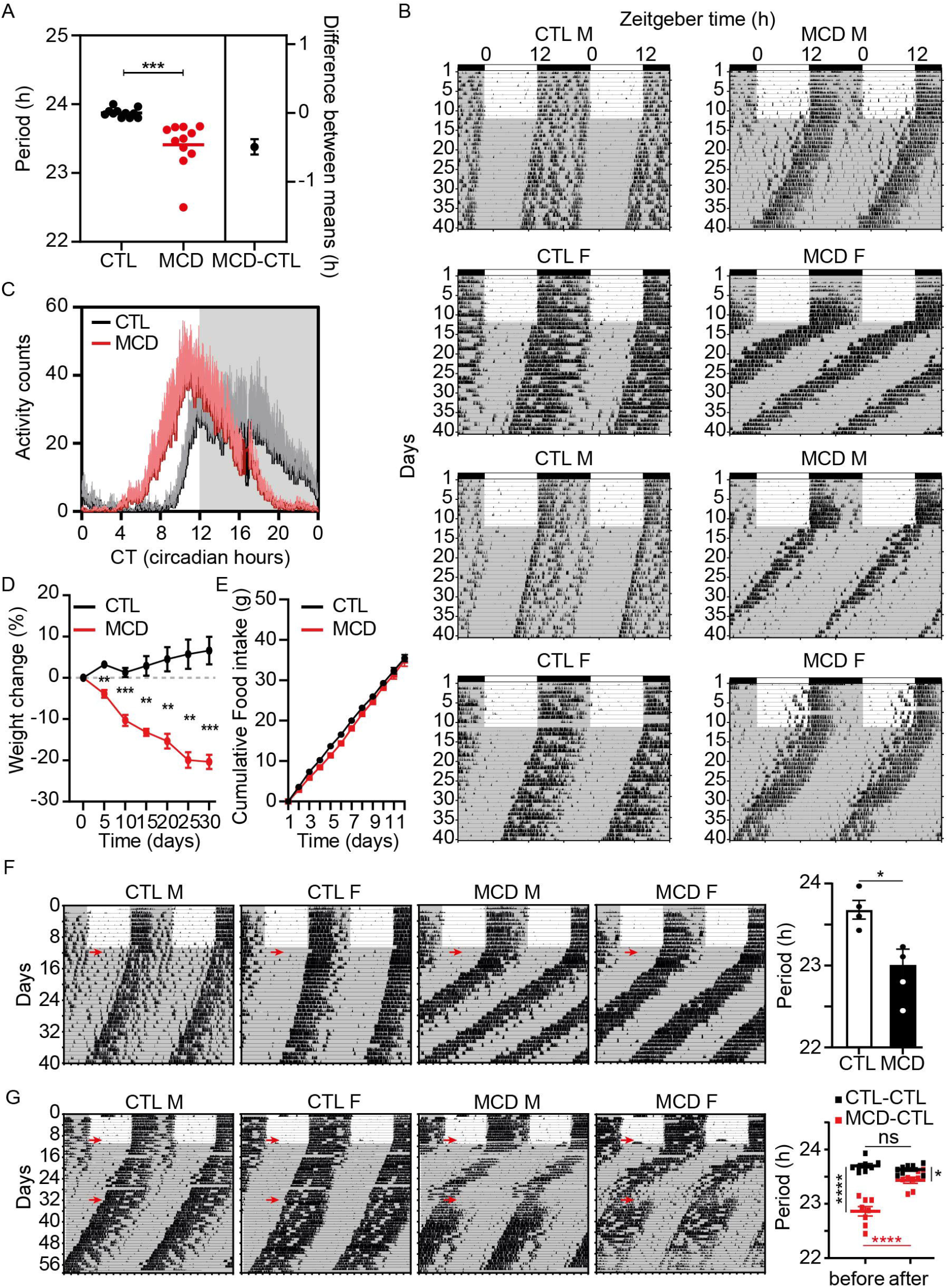
Methionine-choline deficiency affects circadian locomotor activity rhythms. **A**, Circadian locomotor activity rhythm periods of control- or MCD-fed animals analysed by Welch’s t test. The right part of the graph shows the mean period difference (+/- SEM) between the two groups. **B**, Actograms of 2 males and 2 females fed with control (left) or MCD diet (right). Mice were in a 12h/12h light/dark cycle for 12 days before being exposed to constant darkness. Grey background indicates darkness. *Zeigeber* time (X axis) indicates the initial timing of light/dark cycles, lights on at ZT0 and off at ZT12. **C**, Average activity counts +/- SEM (half error bar, above) from mice shown in B; note the earlier activity rise and sharper peak of MCD-fed mice. **D**, Mean relative (% of starting weight +/- SEM) body weight of n = 6 mice (3m + 3f) fed with the control or MCD diet for 30 days, analysed by Two-Way ANOVA (all sources of variations p <0.01) followed by Šídák’s multiple comparisons test. **E**, Cumulative food intake (mean +/- SEM) of n = 4 mice (2m + 2f) fed the control or MCD diet for 12 days, analysed by Two-Way ANOVA (Time, p<0.0001; Diet, p=0.18; Interaction, p=0.79) followed by Šídák’s multiple comparisons test. **F**, Representative actograms of mice fed control or MCD diet starting from the second day in constant darkness, as indicated by a red arrow. Before that time, all mice received to control diet. Right graph shows mean period +/- SEM of n = 4 mice analysed by Welch’s t test. **G**, Representative actograms of mice fed control or MCD diet starting from the last day in a light/dark cycles, as indicate by the first arrow. On the 32^nd^ day of recording, MCD-fed mice were then provided with the control diet until the end of the experiment. Right graph shows mean period +/- SEM of n = 8 mice (4m + 4f) before and after the swap from MCD to CTL diet analysed by Two-Way ANOVA with repeated measures. *, p<0.05; **, p<0.01; ***, p<0.001; ****, p<0.0001.

### Methionine/Choline deficiency reprograms the SCN circadian transcriptome

Given the fact that circadian locomotor activity rhythms are controlled by the suprachiasmatic nucleus of the hypothalamus (SCN), where the master circadian clock resides, changes in circadian behaviour seen in mice fed with the MCD diet in constant darkness should be mirrored at the level of the circadian transcriptome. We thus performed RNASeq on laser-microdissected SCN, from brains sampled every 4 hours from *ad libitum* CTL- or MCD-fed mice for 11 days, the first 10 days in a 12h/12h light/dark cycle, the 11^th^ and last day in constant darkness, when sampling started at Circadian Time 00 (CT00 indicates the beginning of the subjective day, and CT12 the beginning of the subjective night). Circadian transcriptomes in CTL- and MCD-fed mice were compared using DryR, a R-based tool for circadian rhythm comparisons^21^. Out of 15,622 genes considered to be expressed in the SCN (see methods), DryR detected 5410 genes with identical circadian profiles in control and MCD samples, 577 and 486 genes that gained or lost rhythms in MCD samples, respectively, as well as 137 genes that changed circadian profile (Fig. 2A and Supplemental Data 1). The most remarkable of these groups was the genes with changed rhythms: while these genes had two main acrophases in CTL SCN, either at CT19 or CT6-8, their acrophase mainly centred around CT5 in MCD SCN (Fig. 2A). Overrepresentation test with all 1,100 (577 + 486 + 137) differentially expressed genes did not yield significant results (not shown), even when limited to the 157 (68 + 51 + 38) genes among them with a good fit for their respective models, *i.e.* with a Bayesian information criterion weight (BICW) > 0.8. However, the same test with the 38 genes with changed rhythms with a BICW > 0.8 lead to one significant biological process: response to stimulus (GO:0050896), to which 21 of the 38 genes belonged (Supplemental Data 1). Consistent with the DryR analysis, a lot of these 38 rhythmic genes showed earlier rise or fall of gene expression (i.e. phase-advanced), including the clock genes Per1 and Nr1d1; the dual-specificity phosphatases Dusp1 and 4; the thioredoxin interacting protein Txnip; the cold-induced RNA binding proteins Rbm3; the immediate-early gene Junb; the neuropeptides Cholecystokinin (Cck) and Prokineticin (Prok2) involved in the regulation of circadian locomotor activity^22–24^; and acetylcholinesterase (Ache) (Fig. 2B). While this phase advance is consistent with the shorter period of locomotor activity rhythms observed in MCD-fed mice, it is not seen in all rhythmically expressed clock genes, including Arntl and Per2, which were allocated by DryR to the group of genes with identical rhythms in both CTL and MCD conditions (model 4) with a BICW > 0.8 (Fig. 2C). This suggests that the effects of MCD are specific only to certain genes that are part of the transcriptional machinery of the circadian clock and its entrainment, rather than a more general effect on the whole clockwork itself. Indeed, as seen for many of these 38 genes with changed rhythms, the effect of MCD appears more pronounced in the subjective morning.

**Figure 2:**
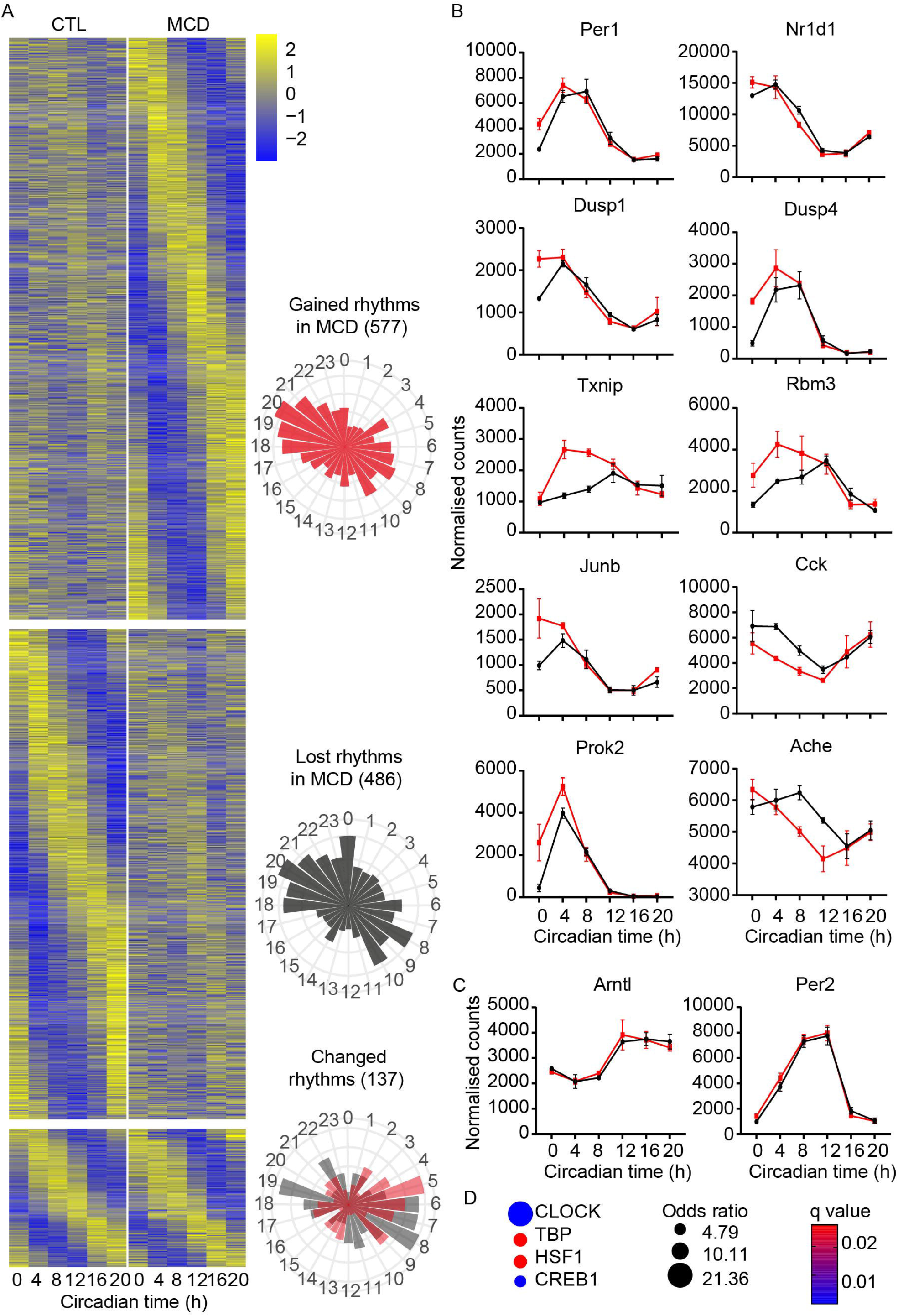
The MCD diet reprograms the SCN circadian transcriptome. **A**, Heatmaps showing profiles of genes with gained, lost or changed rhythms unfiltered by BICW in the SCN of mice fed the MCD diet, compared to mice fed the CTL diet (see also Supplemental data 1). Rayleigh plots on the right show the phase distribution of rhythmic genes. **B**, Profiles of selected representative genes with changed rhythms in the SCN of MCD-fed mice. Data show mean +/- SEM of n=3 animals (2m+1f). **C**, ChiP-X Enrichment Analysis with the 38 genes with changed rhythms (BICW>0.8) as input, showing CLOCK and CREB1 are the most significantly enriched putative transcription factors regulating these genes.

To gain insight into which transcription factors these 38 genes may be regulated by, we use ChiP-X Enrichment Analysis^25^, which revealed the core circadian clock gene CLOCK as the most significant factor, followed by CREB (cAMP Response Element-Binding Protein) (Fig. 2D and Supplemental Data 1). Many of the genes known to be regulated by CREB, including Per1, Nr1d1, Dusp1, Rbm3 and Txnip, have a characteristic higher expression in the morning in the SCN of MCD-fed animals. Together these results suggest that the MCD diet may trigger the cAMP/Ca^++^ signalling cascade in the SCN close to the beginning of the rest phase. This is consistent with the effects of MCD on locomotor activity rhythms in the late night to early morning, when MCD-fed animals show inhibited activity, and in line with high *Per1* and *Prok2*, and generally the activation of cAMP pathway, being associated with low locomotor activity in mice^23,26,27^. Earlier activation of this pathway in the SCN would in turn leads to a phase advance and/or shorter period which potentially explains the effects of MCD on circadian locomotor activity rhythms.

### Methionine/Choline deficiency reprograms the liver circadian transcriptome

The liver has been the main target of studies using the MCD diet. Some evidence that after 4 weeks the MCD diet mildly changes circadian clock gene expression in the liver have been presented^28^, but the effects of methionine/choline deficiency on the liver circadian transcriptome remain unknown. This is a significant knowledge gap, as it is known 10-20% of the liver transcriptome, with many genes coding for rate-limiting enzymes and key regulators of metabolism, is under the control of the circadian clock and feeding/fasting cues^29^, and there should therefore be extensive circadian nutrigenomic interactions with the MCD. We thus analysed the effects of the MCD diet on the circadian hepatic transcriptome, from the same mice used above.

DryR detected a profound reprogramming of the liver circadian transcriptome caused by the MCD diet, with 2650 genes gaining rhythms, 1364 losing rhythms and 1065 with changed rhythms (Fig.3A and Supplemental Data 2). These numbers went down when only genes with a BICW>0.8 were considered (971, 312, 454). Many of the genes losing rhythms were expressed in the early active phase or in the middle of the rest phase in CTL-fed animals. In contrast, the acrophase of genes gaining rhythms was either in the middle of the active phase or at the end of the rest phase in MCD-fed animals. Overrepresentation test with all 1737 genes revealed significant enrichments including negative regulation of ubiquitin-dependent protein catabolic process, positive regulation of TOR signalling, regulation of signal transduction by p53 class mediator, protein-RNA complex assembly and macromolecule methylation, indicating profound effects of the MCD diet on nutrient-dependent regulation of translation and protein folding. (Fig. 3B and Supplemental Data 2).

**Figure 3:**
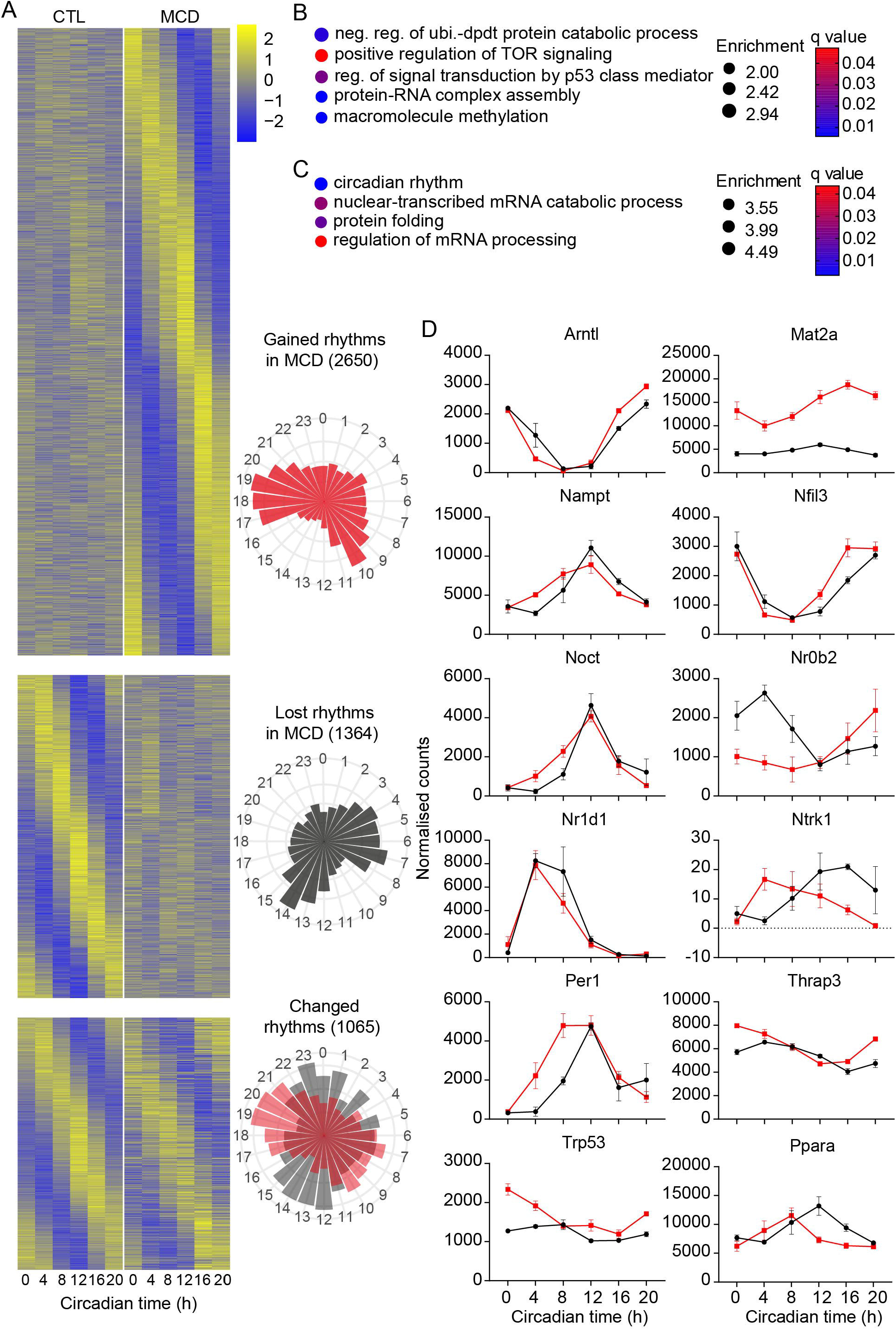
The MCD diet reprograms the liver circadian transcriptome. **A**, Heatmaps showing profiles of genes with gained, lost or changed rhythms unfiltered by BICW in the liver of mice fed the MCD diet, compared to mice fed the CTL diet (see also Supplemental data 2). Rayleigh plots on the right show the phase distribution of rhythmic genes. Notice the difference in gene expression phase between CTL- and MCD-fed mice. **B**, Selected results from overrepresentation analysis with the 1737 genes (BICW>0.8) gaining (model 3) or losing (model 2) rhythms in MCD-fed animals (Supplemental data 2). **C**, Selected results from overrepresentation analysis with the 454 genes (BICW>0.8) with changed rhythms in MCD-fed animals (Supplemental data 2). **D**, Gene expression profiles of selected genes belonging to the circadian rhythm ontology in C. Data show mean +/- SEM of n=3 animals (2m+1f).

A similar phase change of gene expression between CTL- and MCD-fed animals can be seen with the 1065 genes changing expression rhythms (Fig. 3A). Overrepresentation test with the subset of these genes with a BICW>0.8 (454 genes) revealed, perhaps not surprisingly since these genes remain rhythmic in both conditions, significant enrichments in circadian rhythm, nuclear-transcribed mRNA catabolic process, protein folding and regulation of mRNA processing (Fig. 3C and Supplemental Data 2). In the circadian rhythm ontology were the clock genes Arntl, Per1, Nr1d1 and clock-controlled genes Nfil3, Nampt, Noct, Ntrk1, Mat2a, Nr0b2, Thrap3, Trp53 and Ppara, all affected by the MCD diet in different ways but often with a noticeable phase advance or shorter period as observed in the SCN (Fig. 3D). Per1 is especially notable in that regard, in the liver showing a much earlier rise in the morning (Fig. 3D). Together these data show that the MCD diet profoundly affects the circadian clock and its outputs.

One gene in that circadian rhythm ontology, Mat2a, attracted our attention. Not only its mRNA rhythm changed in the liver of MCD-fed mice, but also showed overexpression at all time points (Fig. 3D). Mat2a codes for methionine adenosyltransferase 2a that synthesise S-adenosylmethionine (SAM) from methionine and ATP. It is ubiquitously expressed (unlike its liver-specific homologue Mat1a) but has not been consistently flagged as a gene under the control of the circadian clock. Its inclusion in the circadian rhythms ontology originates from a single 2005 report that its mRNA and protein expression oscillate in the rat pineal gland via a cAMP-dependent mechanism^30^. In our data, *Mat2a* shows a low amplitude rhythm (∼1.5-fold) in CTL liver, but oscillates at ∼3-fold higher baseline, with a higher amplitude (∼2-fold) and a phase delay in the liver of MCD-fed animals (Fig. 3D).

The data on Mat2a prompted us to probe our dataset for other genes involved in 1-carbon metabolism, which we have previously shown is enriched with rhythmically expressed genes in the liver^17^. Out of a list of 49 genes coding for enzymes and transporters involved in 1C metabolism, 27 genes had a BICW>0.8: 7 were classified as non-rhythmic in both conditions (Aldh1l2, Mtap, Ftcd, Sms, Mthfsd, Slc25a26, Mthfd1), 1 lost rhythm in MCD-fed liver (Srm), 4 gained rhythms in MCD-fed liver (Mat2b – the regulatory beta subunit of methionine adenosyltransferase^31,32^, Cth, Chdh, Tyms), 13 had identical rhythms in both condition (Slc19a1, Mtrr, Bhmt2, Slc25a32, Dhfr, Mthfd1l, Bhmt, Ahcy, Fpgs, Amd2, Gart, Tat, Atic) and 2 had changed rhythms (Mat2a, Cbs) (Supplemental Data 2).

Looking closer at the profiles of some of these genes with “identical” or “non-rhythmic” expression rhythms in both conditions, it quickly became obvious that this circadian model classification did not fully capture the effects of MCD in the liver. Indeed, DryR can find rhythmic genes and differences in phase and amplitude between conditions, but has not been designed to detect differences in baseline. Therefore, to complement DryR analysis and detect genes that are significantly up- or down-regulated by the MCD diet, we analysed our data pair-wise (CTL versus MCD) at each time point separately with DESeq2 (Supplemental data 3). This resulted in a list of 7316 significantly-regulated genes in at least one time point. To identify genes and processes affected by the MCD diet independently of their circadian rhythms, we selected genes significant in at least three of the six time points (2,239), which were then used for overrepresentation analysis. This analysis revealed enriched ontologies reflecting the inflammatory condition and profound metabolic changes in the liver of MCD-fed animals (Fig. 4A), including 1-C metabolism, with 16 of the 49 1C-related enzymes (Mthfd1l, Ahcy, Amd2, Mat2a, Apip, Mthfd1, Ahcyl, Mthfd2, Ftcd, Dhfr, Shmt2, Cbs, Aldh1l1, Shmt1, Slc19a1, Aldh1l2) significantly regulated by the MCD diet (Fig 4B, Supplemental Data 3).

**Figure 4:**
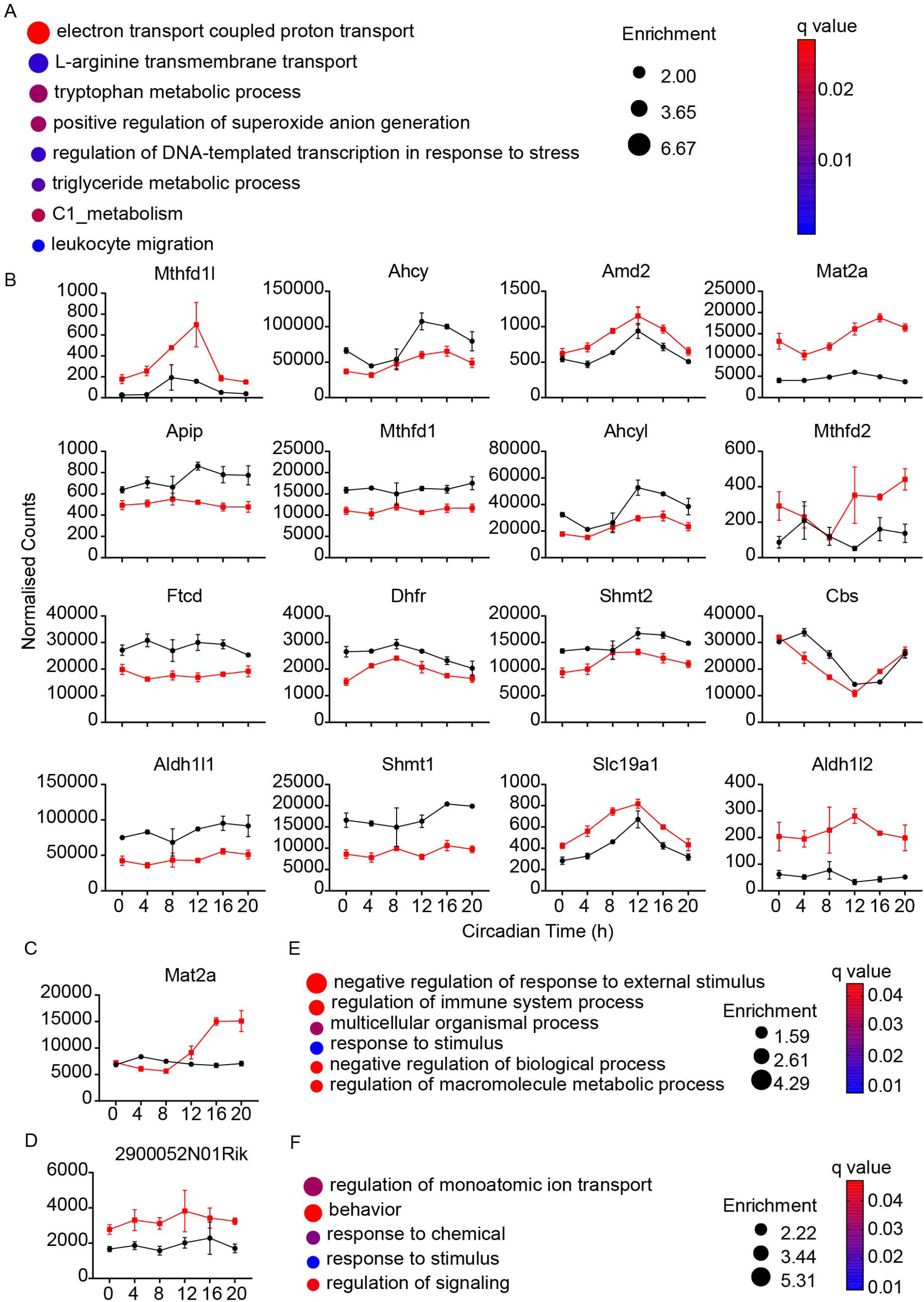
The expression of 1-C metabolism enzymes in the liver and SCN are affected by the MCD diet. **A**, Selected results from overrepresentation analysis with the 2,239 genes significantly regulated by the MCD diet in the liver in at least 3 of the 6 time points (Supplemental data 3). **B**, Gene expression profiles of enzymes involved in 1-C metabolism whose expression is significantly regulated by the MCD diet in the liver in at least 3 of the 6 time points, as detected by DESeq2 (Supplemental data 3). Data show mean +/- SEM of n=3 animals (2m+1f). **C**, *Mat2a* expression in the SCN; data show mean +/- SEM of n=3 animals (2m+1f). **D**, *2900052N01Rik* expression in the SCN; data show mean +/- SEM of n=3 animals (2m+1f). **E**, Selected results from overrepresentation analysis with the 148 genes significantly regulated by the MCD diet in the SCN in at least 1 of the 6 time points (Supplemental data 3). **F**, Selected results from overrepresentation analysis with the 56 genes significantly regulated by the MCD diet in the SCN at CT0 (Supplemental data 3).

### Methionine/Choline deficiency induces circadian oscillations of *Mat2a* in the SCN

We next explored our SCN DryR results for the expression of the same 1C enzymes. 15 genes had a BICW>0.8: 6 genes were identified as non-rhythmic (Dhfr, Aldh1l1, Mthfr, Amd2, Mthfd2l, Amt) and 8 genes had identical rhythms (Aldh7a1, Mtr, Ahcyl2, Mtap, Mthfd2, Slc46a1, Mat2b, Mthfd1l). Interestingly, Mat2a was the only 1C-related gene that gained rhythm in the SCN of MCD-fed animals (Fig. 4C and Supplemental Data 1). Pairwise comparisons by DESeq2 for each time point reinforced our observations made with DryR that the effects of MCD in the SCN are milder and time-specific (Supplemental Data 4). In contrast to the liver, in the SCN there was only one gene significantly regulated by the MCD diet in at least 3 time points: *2900052N01Rik*, a LncRNA of unknown function upregulated by inflammatory injury^33^ (Fig. 4D). There were only 7 genes significantly regulated in at least 2 time points: Noxred1, Rab43, Gm45140 (locus overlapping with Rab43), Gm9294 (pseudogene), Tsc22d3 and Txnip, with Mat2a the only 1C-related gene among them (Supplemental Data 4). Moreover, there were more significantly regulated genes (56) at CT00 than at any other time points (CT04, 12; CT08, 29; CT12, 15; CT16, 20; CT20, 23). Since the list of MCD-regulated genes in the SCN is much smaller than that in the liver, we performed overrepresentation analysis with all 148 genes significantly regulated in at least one time point. This analysis confirmed our previous data with DryR: “Response to Stimulus” was the most significantly enriched ontology (Fig. 4E, Supplemental Data 4). The same analysis with significant genes at CT00 yielded similar results (Fig. 4F, Supplemental Data 4). The effects of the MCD diet in the SCN suggest a specific nutrigenomic link between methionine/choline availability and the regulation of *Mat2a*.

### Methionine/Choline deficiency affects 1-C metabolites in the liver and brain in circadian time-dependent manner

The SCN and liver transcriptome analyses have shown the MCD diet regulates the expression of enzymes involved in 1C metabolism, often in a circadian time-specific manner, which suggest 1C metabolites themselves are affected by the MCD diet. Previous publications have not drawn a consistent picture on the effect of the MCD diet on systemic 1-carbon metabolism. Lower levels of SAM in the liver under MCD have been reported in mice and rats^3,34^ but other authors failed to show any significant changes in hepatic and brain SAM, SAH, choline, betaine and cystathionine^16^. Considering the relationship between 1-carbon metabolism and circadian rhythms^17,35–38^ and the gene expression results reported here, these discrepancies may originate from the failure to control the time of day at which these metabolites were measured.

To define whether 1-carbon metabolism is affected by the MCD diet in a circadian time-dependent manner, we measured selected 1-carbon metabolites in the SCN and liver of mice fed control or MCD diet *ad libitum* for 10 days in a 12h/12h light/dark cycle. On the 11^th^ day, mice were in constant darkness and tissue samples were taken at different time points across 24 hours, either every 4 hours for 24 hours for the SCN, or at circadian time 4 and 16 for the liver. Choline, Methionine, methylthioadenosine (MTA), cystathionine, SAH, SAM and sarcosine had lower levels in liver of MCD-fed animals, but the decrease in cystathionine, methionine and SAH was significantly more pronounced at CT16, during the night when mice are active and eat (Fig. 5A). In contrast, hepatic betaine levels were lower at CT4 but higher at CT16, while serine and glycine levels were indistinguishable between WT and MCD livers at CT4, but higher at CT16 (Fig. 5A).

**Figure 5:**
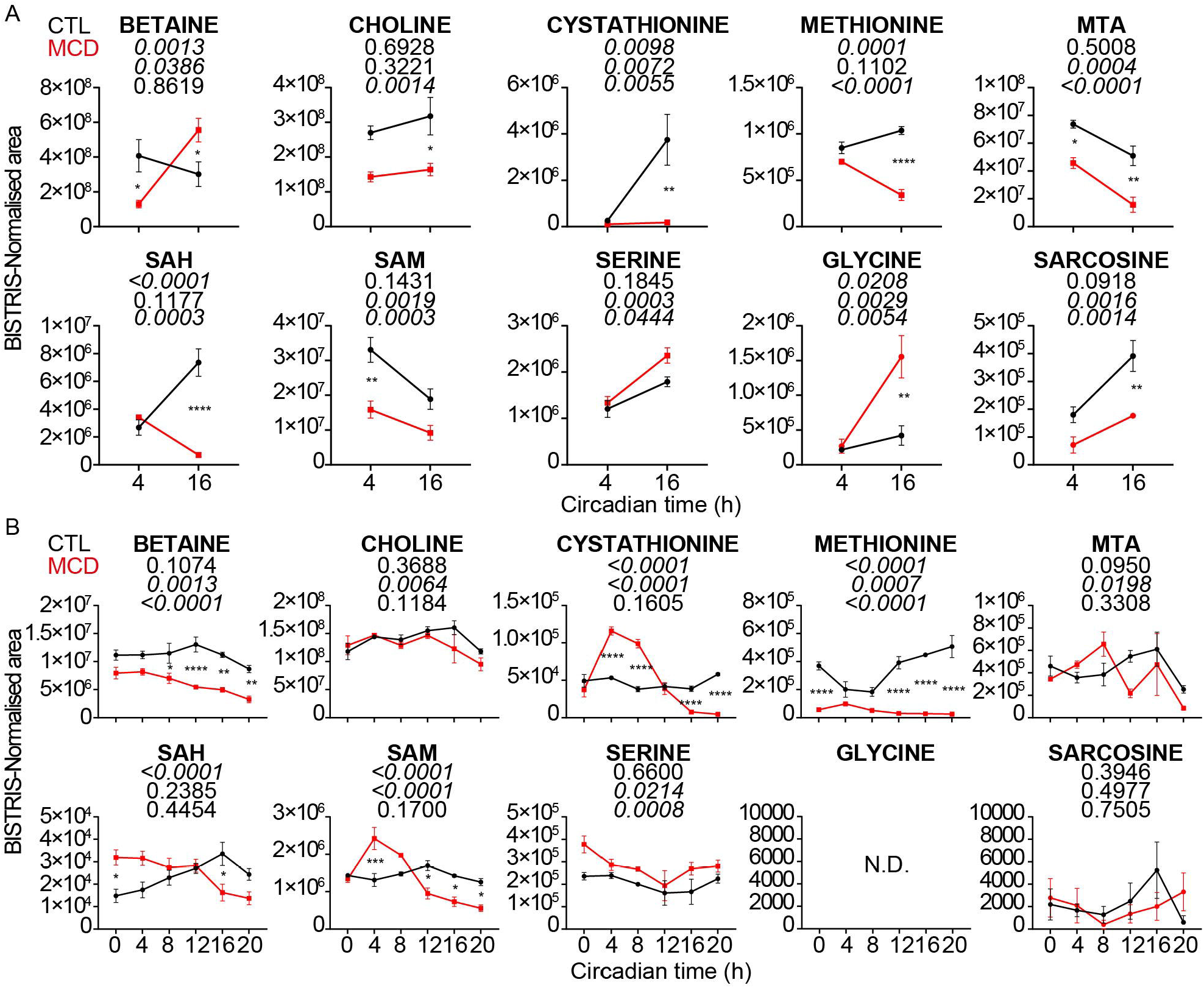
The MCD diet disrupts 1-C metabolism in the liver and SCN in a circadian time-dependant manner. **A**, Metabolite quantification in the liver of control- or MCD-fed animals at CT4 and CT16. Data shows mean +/- SEM of n= 4 mice (2m + 2f), analysed by 2-Way ANOVA followed by Tukey’s multiple comparison test, comparing CTL vs MCD diet for each time point. P values for each source of variation (from top to bottom: interaction, time and diet) are written under each metabolite’s name, italicized when significant (<0.05). **B**, Metabolite quantification in the SCN of control- or MCD-fed animals across 24 hours. Data show mean +/- SEM of n= 4 mice (2m + 2f), analysed by 2-Way ANOVA followed by Tukey’s multiple comparison test, comparing CTL vs MCD diet for each time point. P values for each source of variation (from top to bottom: interaction, time and diet) are written under each metabolite’s name, italicized when significant (<0.05).

In the SCN, SAM and SAH were lower during the active phase under MCD, but higher than control during the day (Fig. 5B). Methionine was significantly lower in SCN samples under MCD at all time points, but displayed antiphasic rhythms of similar amplitudes under both conditions, so that the most pronounced differences in methionine levels between diets was observed towards the end of the night, at CT20 (Fig 5B). Cystathionine showed little circadian variations in CTL conditions, but a high-amplitude rhythm with high daytime levels was observed in the SCN of MCD-fed mice (Fig 5B). Choline showed weak but significant 24h variations, and was not significantly affected by the diet (Fig 5B). Betaine was lower in the SCN of MCD-fed animals at all time points, especially at night (Fig 5B). MTA levels showed daily fluctuations in both control- and MCD-fed animals (Fig 5B).

Together these data demonstrate that the MCD diet has pronounced systemic effects on 1-C metabolism, but that the direction and magnitude of these changes are dependent on circadian time, likely related to feeding/fasting rhythms.

### Methionine/Choline deficiency affects splicing in the liver and the SCN

Our data so far have shown that the expression of key 1-C enzymes and their substrates are regulated by the MCD diet, and *Mat2a* has been highlighted as one of the most strongly regulated genes in both the liver and the SCN. Intriguingly, it has been shown that SAM, via the *N*^6^-adenosine methylation of *Mat2a* mRNA by the methyltransferase METTL16, regulate the alternative splicing of *Mat2a*^39–41^. When SAM is abundant, methylation of *Mat2a* mRNA leads to intron retention, accelerated *Mat2a* mRNA degradation and decreased MAT2A expression. In contrast, when SAM is limiting, decreased *Mat2a* methylation leads to increased expression of the coding transcript and MAT2A translation, to synthesise more SAM. To our knowledge, conditions where such posttranscriptional *Mat2a* regulation operate have not been shown in adult mammals *in vivo*. Our data therefore provide an opportunity to define whether intron retention within the *Mat2a* transcript is regulated by the MCD diet and circadian time, and to identify new potential genes that may be regulated in the same way.

We first probed our RNASeq data for significant changes in transcriptome-wide exon usage in the liver between CTL and MCD-fed animals at each time point (Supplemental data 5). A total of 3613 significant events corresponding to 2272 genes were identified, with 127 significant events in at least 3 of the 6 time points. Only 4 splicing events were significantly affected by the MCD diet in all of the 6 time points: 2 exons in *Scd1*, and 2 exons in *Mat2a* (Supplemental data 5). Four sequential exons of *Mat2a*, E006-009, were found to be differentially spliced in at least 4 of the 6 time points: these exons in the gene model correspond to the intron 8-9 from transcript ENSMUST00000059472.10 that is retained in transcript ENSMUST00000205335.2 (Fig. 6A and Supplemental data 5). In the liver of MCD fed mice, the usage of these 4 exons was consistently lower (*i.e.* negative MCD/CTL log2 fold ratio) than in CTL-fed mice, but variations in the differential usage of these exons were also observed between time points. The most negative ratio (∼-3) was seen at CT20, when the difference in *Mat2a* expression between CTL- and MCD-fed animals was most pronounced (Fig. 6B and Supplemental data 5), which may be related to changes in the concentration of 1C metabolites including methionine, SAM and SAH (Fig. 5A). Two other exons, E019 and E023, also corresponding to intronic regions in the 2 canonical transcripts ENSMUST00000059472.10 and ENSMUST00000205335.2, were significantly affected by the MCD diet at CT04 and CT20, their usage decreased with the MCD diet (Fig. 6B and Supplemental data 5). Together these data show that *Mat2a* splicing is regulated by methionine/choline availability in the liver.

**Figure 6:**
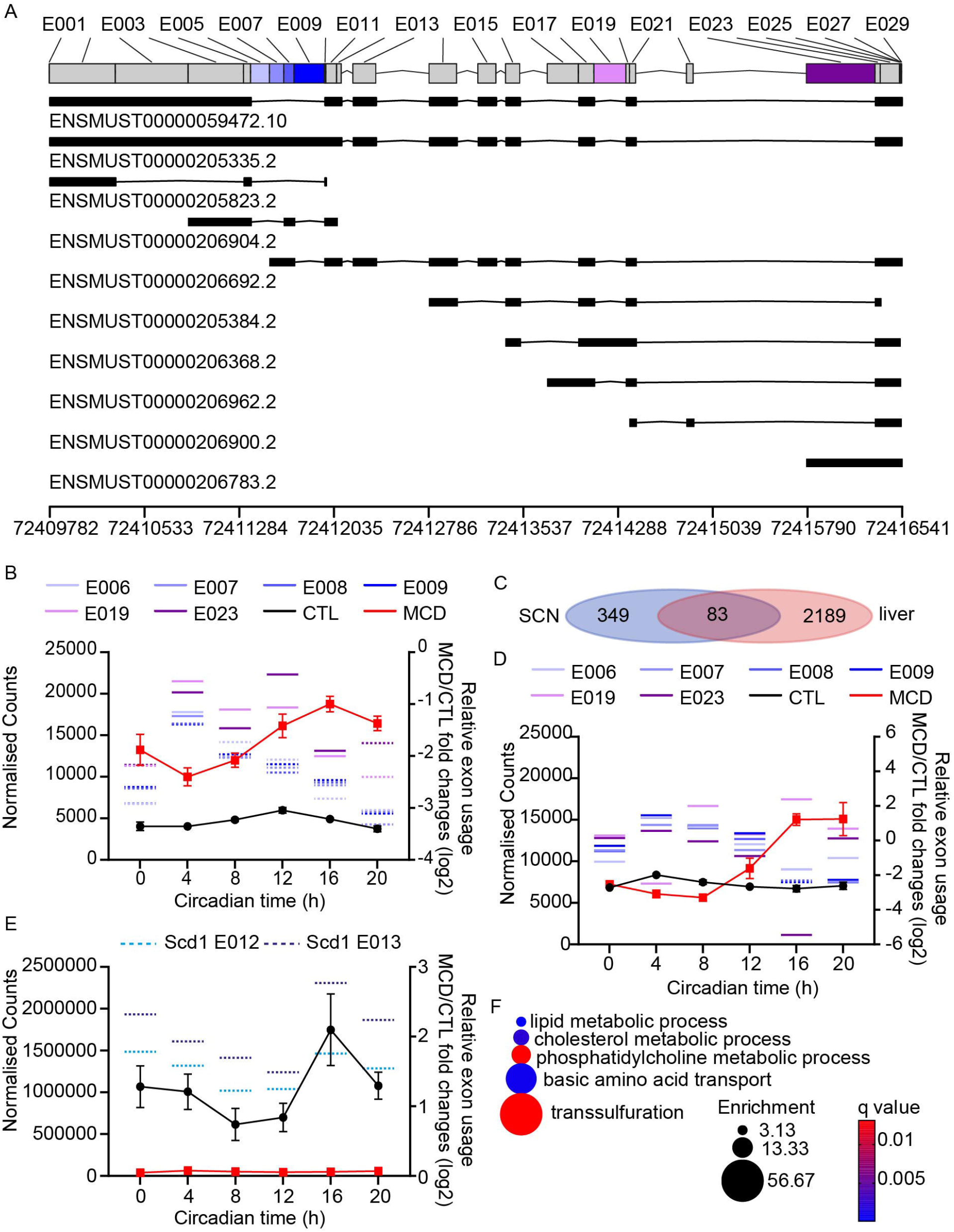
The MCD diet triggers changes in alternative splicing. **A**, Representation of the Mat2a gene model on chromosome 6 and its various transcripts. The gene model shown at the top is a flattened representation of all potential exons, represented as grey or coloured boxes, separated by intronic sequences shown as think black lines. Exons are counted from the 5’ end of the plus strand, but since Mat2a is encoded in the minus strand, the 5’ end of *Mat2a* is on the right, starting with exon 29. Transcript ENSEMBL IDs are shown under each transcript representation, and genomic locations are shown at the bottom. Coloured exons are those whose usage has been found to be significantly regulated by the MCD diet, data shown in B and D. **B**, Mean usage of *Mat2a* exon 6-9, 19 and 23, plotted on the right axis together with *Mat2a* expression from Fig. 3D and the left; liver data. Dotted lines indicate CTL vs. MCD exon usage significance (q<0.05) in DEXSeq analysis (Supplemental data 5). **C**, Venn diagram showing the numbers of genes with significant exon usage in the SCN and the liver from DEXSeq analysis. **D**, Mean usage of *Mat2a* exon 6-9, 19 and 23, plotted on the right axis together with *Mat2a* expression from Fig. 3D on the left; SCN data. Dotted lines indicate CTL vs. MCD exon usage significance (q<0.05) in DEXSeq analysis (Supplemental data 6). **E**, Usage of *Scd1* exon 12 and 13, plotted on the right axis together with *Scd1* expression from Supplemental data 1; liver data. Dotted lines indicate CTL vs. MCD exon usage significance (q<0.05) in DEXSeq analysis (Supplemental data 5). **F**, Selected results from overrepresentation analysis with the 127 genes with significant splicing events in at least 3 of the 6 time points in the liver (Supplemental data 5).

Next, the same exon usage analysis was performed with the SCN data. As observed for the gene expression analysis, the effects of MCD were more subdued in the SCN. 467 events corresponding to 432 genes were significant in at least 1 time point, and only three events (Septin4, Srp54b, Ccdc177) were significant in two of the six time points (Supplemental data 6). Mat2a was among the 83 genes co-regulated in the liver and the SCN (Fig. 6C). The usage of exons E008 and E009 was significantly lower at CT16 in the SCN of MCD-fed animals, when *Mat2a* expression increases (Fig. 6D and Supplemental data 6) and in line with lower methionine and SAM levels (Fig. 5B), indicating that the splicing of *Mat2a* is also regulated in the SCN by methionine and choline availability.

Mat2a was not the only gene whose splicing was regulated by methionine/choline availability, especially in the liver. *Scd1* was the only other transcript showing significant splicing events at all time points. The physiological functions of *Scd1* splicing in the liver is not known but it is clear that the usage of the two 5’-most exons of this gene, E012 and E013, is higher in the liver of MCD animals (*i.e.* positive MCD/CTL log2 fold ratio), likely leading to a truncated transcript and premature degradation, judging by the ∼20-fold lower levels of *Scd1* in MCD-fed liver (Fig. 6E). *Scd1* codes for stearoyl-Coenzyme A desaturase 1, a lipogenic enzyme involved in monounsaturated fatty acids synthesis^42^, and splicing of *Scd1* may have a critical role in the development of steatosis and weight loss caused by the MCD diet. Indeed, Enrichment analysis with alternative splicing events significant in at least 3 time points in the liver (127 genes) revealed notable enrichments in lipid and cholesterol metabolism, phosphatidylcholine metabolism, amino acid transport and transsfufluration, all directly related to methionine and choline metabolism (Fig. 6F and Supplemental data 5). These data also provide evidence that the splicing of many other transcripts, including the neutral amino acid transporter *Slc38a4*, the Dihydropyrimidinase *Dpys*, the 1C-related enzyme *cystathionine beta-synthase* (*Cbs*), the *Scd1* homologue *Scd2*, is regulated by methionine/choline availability in the liver (Supplemental data 5).

## Discussion

The potent effects of the MCD diet on circadian behaviour is surprising, as the SCN has been shown to be relatively resistant to stress and peripheral hormones^43,44^. High-fat diets cause a progressive but mild lengthening of the circadian period^45^, and evidence for an effect of high-fat diet on circadian gene expression in the SCN are limited to several clock genes measured by real-time PCR, *in situ* hybridisation or via luciferase reporters, data not altogether consistent showing the molecular clock in the SCN appears little affected by high-fat diets^45–47^. Similarly, restricted feeding to the light phase changes circadian oscillations in the mouse liver but not the SCN^48–50^. Here as well, overall the circadian expression of core clock and clock-regulated genes seems little affected. Yet, circadian behaviour responded quickly to a change in diet, either from control to MCD or *vice versa*: within a few days phase advances (CTL→MCD) or delays (MCD→CTL) are observed, with the circadian period rapidly adjusting (Fig. 1F). Moreover, compared to control-fed animals, animals fed the MCD diet often showed noisy activity onsets in constant darkness (Fig. 1). This could be due to a lack of robustness leading to unstable period length and/or to phase advances of variable magnitude occurring every day due to MCD diet intake at night. Considering our data, we favour the latter, as it may appear consistent with the earlier rise in cAMP/Ca^++^-responsive genes in the SCN of MCD-fed animals in the morning. This is similar to the phase-advancing effects of timed-hypocaloric feeding in mice^45^ and the short circadian period of long term food-restricted rats^51^. More work will be required to understand the underlying mechanisms.

A recent report showed that the lack of methionine or other essential amino acids failed to affect circadian locomotor activity rhythm period in C57BL/6 mice^52^, suggesting that the lack of both methionine and choline was necessary to elicit the observed effects on the circadian period reported here. Other differences between the two reports is the use of Teklad’s diet TD.140119 and control TD.01084^52^, while we used our own formulations (TD.220158 and its methionine/choline replete control TD.220157) that contain half the soybean oil content to limit steatohepatitis. TD.140119 contains 4.95 g/Kg serine, compared to 3.5 in TD.01084 and in both our diets. Since serine is a 1-carbon donor for homocysteine remethylation to methionine via 5-methyltetrahydrofolate, it is possible that the higher serine content in TD.140119 may have compensated for the lack of methionine. Further work should clarify the discrepancies between the two reports. In our two diets, to keep the same total weight constant we use sucrose to compensate for the lack of methionine and choline: TD.220158, sucrose 355.88 g/Kg; TD.220157, sucrose 344.98 g/Kg; but these diets are nevertheless isocaloric due to carbohydrate/protein balance (3.7 Kcal/g). There was no significant difference in food intake between control- and MCD-fed mice (Fig. 1E).

As shown in our analyses, DryR is very sensitive to differences in period, phase and amplitude in cycling gene expression between groups, but at the same time blind to changes in baseline, so that it cannot detect genes that are up- or down-regulated with a similar magnitude at every time point. This is very common among circadian omics analysis packages. Another R package, CompareRhythms, is different from DryR as it is based on hypothesis testing to define genes that have significantly different amplitude or phase, while DryR performs selection of harmonic regression model fits to different rhythm categories^21,53^. While compareRhythms also ignores potential differences in baseline, it has been argued that it is less sensitive to false positives hits than methods such as DryR. In addition, compareRhythms also applies an amplitude threshold —albeit arbitrary— for a gene to be considered rhythmic. We thus decided to analyse our RNASeq data from the liver and the SCN by compareRhythms (Supplemental data 7). In the liver, a total of 1455 genes were detected as significantly differentially rhythmic by CompareRhythms, but there was relatively little overlap between these genes and the 1737 genes with BICW>0.8 that lost, gained or had changed rhythms identified by DryR (Fig. 7A). Decreasing the BICW threshold for DryR lead to more genes identified as differentially rhythmic by compareRhythms to be also detected by DryR, but at the same time increased the fraction of genes detected only by DryR (Fig. 7A). Describing these latter genes (at least for those with a BICW>0.8) as false positives would be inaccurate, judging by a few of them (Fig. 7B). Likewise, the 30 genes detected only by compareRhythms seem to show some effects of the MCD diet (Fig. 7C). Such poor overlap between rhythm detection methods and the limitations to comparing them have already been flagged elsewhere^54,55^. Regardless of the methods used and the arbitrary thresholds applied, however, it is clear that the MCD diet exert profound effects on the hepatic transcriptome. Analysis of the SCN data with CompareRhythms lead to only 6 differentially rhythmic genes to be detected: Mat2a, Txnip, Per1, Mmp14, Tsc22d3 and Dnajb1 (Fig. 7D and Supplemental data 7). These genes were also detected by DryR, either with changed rhythms (Txnip, Per1, Mmp14, Tsc22d3 and Dnajb1) or gain of for Mat2a (BICW>0.8). Mat2a was the most significant gene (FDR = 3.22E-22) in the compareRhythms analysis, further highlighting it as a highly specific response of the 1C metabolism in the SCN to the MCD diet. While DryR and CompareRhythms may each have their advantages, as we illustrate here overlooking differences in baseline limits the insights gained from the comparison of circadian transcriptome datasets between conditions. We would strongly encourage the circadian community to consider this. In this regard, less commonly used packages that use linear models to fit time-dependant changes (circadian or otherwise) in gene expression and able to detect changes in baseline, such as LimoRhyde^56^ and MaSigPro^57,58^, may be useful.

**Figure 7:**
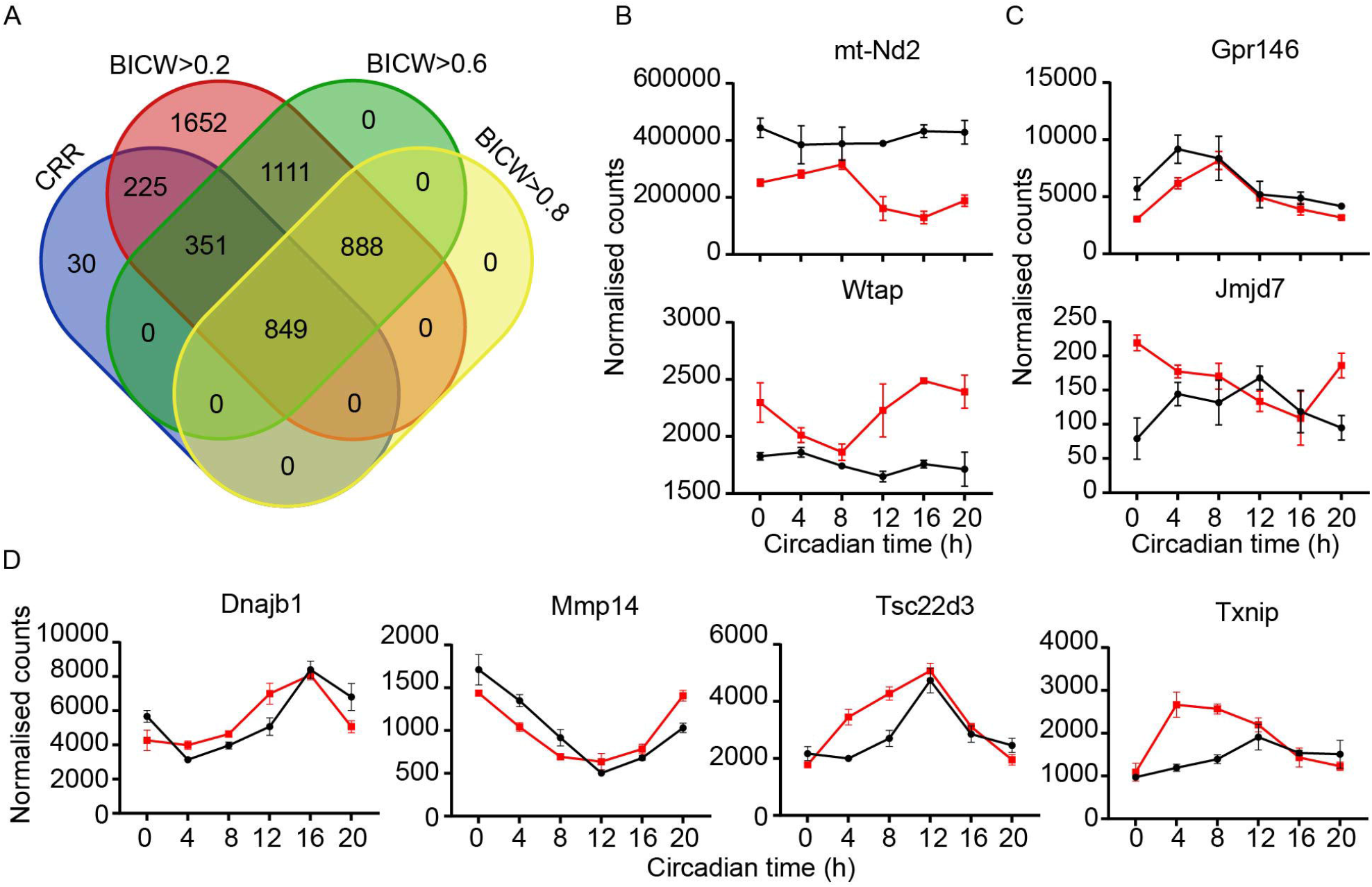
Differences between DryR and CompareRhythms outputs reveal the complexity of comparing circadian omics datasets. **A**, Venn diagram showing the number of unique genes flagged as significant by CompareRhythms (CRR) and DryR using different BICW thresholds. **B**, Gene expression profiles of selected genes identified only by DryR (BICW>0.8) as significant. **C**, Gene expression profiles of selected genes identified only by CRR as significant. **D**, Gene expression profiles of the only four other genes flagged as significant by CRR in the SCN, besides Mat2a and Per1. Gene expression data show mean +/- SEM of n=3 animals (2m+1f).

A physiological meaning behind the effects of MCD on circadian behaviour may be proposed: the absence of methionine and choline from the food is detected several hours into the feeding phase by an unidentified sensor, either of methionine and/or choline themselves or of related metabolites such as SAM or SAH, which triggers first a premature rest, maybe to conserve energy, then an earlier rise (i.e. shorter circadian period) the next day in an attempt to gain a lead in foraging. Since the animal fail to acquire the missing nutrients, the same pattern of rest/activity repeats every day. How this operate remains to be investigated but may be dependent on inflammatory messengers from the liver to the brain, on mTOR-related sensors of methionine and/or SAM including BMT2^59^, and/or on changes in epigenetic and epitranscritomic methyl marks.

## Methods

### Animal experiments

Animal experiments were licensed under the Animals (Scientific Procedures) Act of 1986 (UK) and were approved by the animal welfare committees at the University of Manchester. Eight- to twelve-weeks old C57BL/6J male mice were purchased from Charles River (UK) and acclimatized to the local animal unit for 1 week before the experiment. Mice were then single-housed under a 12:12 h light/dark cycle at ∼350 lux during the light phase and 0 lux during the dark phase for 10-12 days before being exposed to constant darkness (DD) for the remainder of the experiment. Ambient temperature was kept at 21 ± 5 °C, the humidity was ∼51 ± 8%, with water available *ad libitum*. MCD diet TD.220158 or its isocaloric control TD.220157 were provided *ad libitum*, the timing as indicated for each experiment. Throughout the experiment, mice had unrestricted access to a running wheel connected to a wireless transponder allowing real-time recording of activity (Clocklab, Actimetrics), and circadian locomotor activity rhythms period was measured from the actogram by *Chi* square periodogram (Clocklab, Actimetrics).

### RNA extraction

At the appropriate circadian time and under dim red light, mice were killed humanely by cervical dislocation followed by cessation of circulation. Eyes were removed, then under normal room lighting liver (∼40 mg from the centre of the left lobe) and whole brain were immediately dissected and flash frozen in liquid nitrogen or on dry ice, respectively. RNA was extracted form liver samples using standard Trizol procedure (Invitrogen). Brains were cryosectionned (Leica CM3050, head at - 15°C chamber -20°C), first by coarsely trimming the anterior brain until the start of the SCN, then by cutting 23 sequential 25-micrometer coronal sections, each immediately mounted onto a single PEN Membrane Glass Slide (LCM0522, ThermoFisher) and allowed to dry for a few seconds. Slides with mounted sections were warmed at room temperature for a few seconds, then fixed for 3 min in ethanol:acetic acid (19:1), rinsed briefly by immersion in water, stained by immersion in water with 0.2% toluidine blue followed by two washes by immersion in water. All solutions were ice-cold and RNase-free. After the last wash, excess water was flicked off from the slide and slide was placed (sections face up!) onto a plate heater at 50°C for a few minutes until dry. On the same day, SCN were microdissected using a laser dissection microscope (Leica LMD7000; 10× magnification), using 350 microliter RLT buffer with 1% b-mercaptoethanol as a collection/lysis buffer (RNeasy micro kit, Qiagen). RNA was extracted following manufacturer’s instructions (RNeasy micro kit, Qiagen).

### Metabolites extraction

Liver and brain tissues were dissected and flash frozen as described above. For liver tissues, 800µl of 50:50 water:methanol, 125 ng/ml BIS-TRIS solution were added per 50mg of tissue in 2ml Safe-Lock tubes (Eppendorf); one 3mm Tungsten carbide bead (69997, Qiagen) was added to each tube. Samples were homogenised for 10 min at 25 Hertz in a TissueLyser II (Qiagen), then centrifiuged at 12,000xg for 10min. Supernatant was transferred to a new tube and kept at -20°C until analysis.

For SCN samples, brains were trimmed in the cryostat (cryostat head at -12°C, chamber -15°C) as described above but then, two sequential 250 microm slices were sectionned, and the SCN was punched out of the two slices and immediately transferred to a pre-frozen 1.5 ml microtube (Eppendorf) kept on dryice. 50 microl of 50:50 water:methanol, 125 ng/ml BIS-TRIS solution precooled close to -20°C were added to the tube that was then vortexed and spun down. Samples were then homogenised in a bath sonicator (Bioruptor, Diagenode) for 5 min, set to high with 30s/30s on/off cycles. Samples were centrifuged at 12,000xg for 10min. Supernatant was transferred to a new tube and kept at -20°C until analysis.

Metabolite quantification by LC-MS/MS was performed by our BioMS technology platform as previously described^13^.

### RNAseq

Total RNA samples were submitted to our Genomic Technologies Core Facility (GTCF) and libraries were generated as previously described^13^. The Galaxy server^60^ was used for all pipelines excepted DryR and CompareRhythms. Raw reads were QCed and trimmed using Trimmomatic^61^ (Sliding window trimming, 4 bases average, average quality required 20) before being aligned to the mouse genome (Release M33 GRCm39) using HISAT2^62,63^ using default options. Transcripts were assembled and counted using Stringtie^63,64^ with GENCODE annotations for M33 (GRCm39), using default options and keeping all annotated genes including all predicted genes, pseudogenes and non-coding RNAs (for a total of 56,884 “genes”). Gene counts output from Stringtie were then used as input from DryR or compareRhythms, pre-filtered based on Cook’s distance and the mean of normalized counts by DESEq2 to remove genes with too low expression values and high residuals (total number of genes from liver, 22,823; SCN, 15,759). DryR assigns genes to 5 separate models, in our case: 1, genes arrhythmic in both conditions; 2, genes losing rhythms in MCD samples; 3, genes gaining rhythms in MCD samples; 4, genes with identical rhythms in both conditions; 5, genes with changed rhythms between the 2 conditions. To only keep genes best fitting these DryR models, a Bayesian information criterion weight (BICW) cut-off of at least 0.8 was applied before downstream analyses when specified.

Stringtie gene counts tables were also used for DESEq2^65^ pairwise comparisons of transcriptome signatures between MCD- and control-fed animals at each time points, also applying DESEq2’s filters. For overrepresentation analyses with these DEseq2 results from the liver, we selected genes significant (FDR<0.05) in at least 3 of the 6 times points, and the background list of genes was composed of all genes (20,804) that had passed DESeq2’s standard filtering in at least one time point. For the SCN samples, we selected all genes significant in at least one time point, and the background genes included all genes (30,068) that had passed DESeq2’s standard filtering in at least one time point.

DEXseq was used for differential exon usage analysis^66^. First, exons were counted from the bam files output by HISAT2 using GENCODE annotations for M33 (GRCm39) prepared for counting by the DEXseq-Count function. Differential exon usage was then analysed from count tables. For each gene, DEXSeq numbers all exons sequentially from all known and predicted transcripts and always from the 5’ end of the reference strand, even for genes that are on the opposite strand.

Overrepresentation analyses were performed using GOEnrichment (https://github.com/DanFaria/GOEnrichment) on Galaxy, using updated ontology (obo) and annotation (gaf) files from geneontology.org, modified with an additional “C1_metabolism” ontology (GO:9999999) as previously described^17^.

## Data availability

Raw RNASeq data, gene counts and exon counts for each sample are available at the NCBI’s Gene Expression Omnibus, series GSE252364. Raw data and actograms for each animal are available upon request.

## Acknowledgements

This work was supported by the Medical Research Council (Future Leaders Fellowship MR/S031812/1) and by the University of Manchester, Faculty of Biology, Medicine and Health Platform Sciences. The authors wish to thank Dr Peter Briggs (University of Manchester, Faculty of Biology, Medicine and Health Platform Sciences) for his excellent support with the local Galaxy service, and the staff at the Genomic Technologies Core facility of the University of Manchester, Faculty of Biology, Medicine and Health. We also thank Prof. Hitoshi Okamura from Kyoto University for providing funding for a pilot MCD experiment at the Department of Systems Biology.

## Contributions

J.-M. F. conceived and designed the study. J.-M. F. and B. Saer. performed the experiments. G. T. performed the metabolite quantifications. A. H. provided advice on RNAseq experimental designs. B. A. performed CompareRhythms analysis. J.-M. F. wrote the first draft of the manuscript, with input from B. Saer. and B. S.

